# The first complete mitogenome of the endangered spotted eagle ray, *Aetobatus ocellatus*, reinforces its classification within the family Aetobatidae

**DOI:** 10.1101/2025.01.13.632509

**Authors:** Alan Marín

## Abstract

The complete mitochondrial genome of the endangered spotted eagle ray, *Aetobatus ocellatus*, was determined for the first time using Next Generation Sequencing (NGS) reads mined from the Sequence Read Archive (SRA) of the GenBank database (BioSample SAMN31811701, collection site: Bohol Island, Philippines). The mitogenome of the spotted eagle ray (GenBank accession BK072016) is 20,217 bp in length and exhibits a typical vertebrate mitogenome organization, consisting of 13 protein-coding genes, 22 tRNA genes, 2 rRNA genes, and a control region. A phylogenetic analysis based on the nucleotide sequences of 13 protein-coding genes from different related species confirmed the species status and classification of *A. ocellatus* as an Indo-West Pacific species. This species was formerly described as *A. narinari*, which is now recognized as being restricted to the Atlantic Ocean. The findings of this study provide significant insights into the complex taxonomic and phylogenetic relationships of Aetobatidae. They may further assist in conservation strategies and help resolve taxonomic uncertainties concerning the classification of species within Aetobatidae, which remains under debate.

## Introduction

Eagle rays of the genus *Aetobatus* (Myliobatiformes: Aetobatidae) are large pelagic fish found in tropical and subtropical seas around the world (White and Last 2016). They face significant threats from strong overfishing and have a slow recovery from overexploitation due to their low regeneration rate, slow growth, and late maturity (Yamaguchi et al. 2021). As a result, their populations continue to decline throughout their distribution ranges (Dulvy et al. 2021; Finucci et al. 2024), leading to all members of this genus being classified as Endangered or Vulnerable on the IUCN red list (IUCN 2024).

The classification of *Aetobatus* species is particularly susceptible to taxonomic confusion due to the interspecific overlap of their morphological features (White et al. 2013). Until 2010, only two nominal species, *A. narinari* and *A. flagellum*, were recognized as valid (White et al. 2010). However, in the past decade, several taxonomic changes have been made with the help of genetic markers. These changes include the resurrection of *A. ocellatus* as the valid Indo-West Pacific species within the *A*. narinari complex; the redescription of *A. flagellum* (White and Moore 2013); the identification of a new cryptic species, *A. narutobiei* (White et al. 2013); and most recently, the resurrection of the family Aetobatidae (White and Naylor 2016).

These taxonomic revisions have led to the recognition of five currently accepted nominal species: *A. flagellum* (Indo-West Pacific), *A. narutobiei* (North-West Pacific), *A. narinari* (Western-Eastern Atlantic), *A. laticeps* (Eastern Pacific), and *A. ocellatus* (Indo-Pacific) (White and Last 2016; Froese and Pauly 2025). Molecular and morphological analysis have confirmed that the last three species belong to the *A. narinari* complex, formerly considered as a monotypic species with a circumtropical distribution (White and Moore 2013). However, it is likely that additional species are part of the *A. narinari* complex and thereby further extensive taxonomic revisions and molecular analyses are still needed to clarify the taxonomy of this complex (White et al. 2013; White and Last 2016; Sales et al. 2019).

Clarifying the taxonomic status of highly exploited and endangered species is crucial for effective stock management and conservation efforts (Pickett et al. 2020). In this context, extensive genomic data and whole mitochondrial genome sequencing serve as powerful molecular tools that have proven to be highly effective in elucidating the phylogenetic relationships among various fish species complexes (Miya and Nishida 2015). Despite the endangered status and complicated taxonomic history of *Aetobatus*, to date only one complete mitogenome have been sequenced for *A. flagellum*, which measures 20,201 bp (GenBank accession NC_022837). Two additional mitogenome sequences for *A. narinari* (15,714 bp; GenBank accession KX151649) and *A. ocellatus* (11,901 bp; GenBank accession JN184054) are available in GenBank. The mitogenome sequence for *A. narinari* is nearly complete, missing only the control region. In contrast, the mitogenome sequence for *A. ocellatus* is incomplete, lacking 25 genes. Consequently, the sequence for *A. ocellatus* has limited utility for robust phylogeny analysis.

To further support the taxonomic status of *Aetobatus* species and enhance understanding of the global phylogeny within this complex, there is a need for the complete determination and analysis of additional mitogenomes. In this study, I present the first complete mitochondrial genome of the endangered spotted eagle ray, *A. ocellatus*, which was assembled from publicly available short genomic reads in the GenBank database. The structure and sequence content of the mitogenome were analyzed to investigate its phylogenetic relationship within the order Myliobatiformes.

## Materials and methods

### Genomic NGS reads mining, mitogenome assembly, and gene annotation

The short genomic reads used to assemble the mitochondrial genome of *A. ocellatus* were obtained from BioSample SAMN31811701, which was sourced from the GenBank database.

According to the BioSample metadata, the specimen of *A. ocellatus* (initially identified as *A. narinari* by the original submitters) was collected from Bohol Island in the Philippines and sequenced by the Smithsonian National Museum of Natural History as part of their “NOAA Genome Skimming of Marine animals” project (BioProject accession: PRJNA720393).

Paired-end FASTQ genomic reads, totaling 2.2 Gbp within 14.6 million reads, were downloaded from the public GenBank SRA repository (SRA accession: SRS15852991). The reads were first trimmed using BBDuk, resulting in 14.5 million remaining reads, and then merged with BBMerge, yielding 4.1 million reads. This process was carried out using Geneious Prime v. 2021.1.1 (Biomatters Ltd., Auckland, New Zealand). Both unmerged and merged reads were independently assembled against the only complete mitogenome reference sequence of *Aetobatus* available prior to this study (*A. flagellum*, 20,201 bp, GenBank accession NC_022837). The assembly employed the “Map to Reference” tool in Geneious Prime with the recommended default parameter settings. The consensus sequences obtained from both unmerged and merged reads were then extracted and manually verified using the alignment tool in Genious Prime.

Protein-coding (PCG), transfer RNA (tRNA), and ribosomal RNA (rRNA) genes were annotated using the “Annotate and Predict” feature in Geneious Prime. This process involved comparing the novel mitogenome of *A. ocellatus* to the complete mitogenomes of closely related species. The positions of the start and stop codons for the PCGs were verified using the Open Reading Frame Finder tool (available at https://www.ncbi.nlm.nih.gov/orffinder/) applying the vertebrate mitochondrial genetic code. The locations of the tRNAs were confirmed using tRNAscan-SE (Chan and Lowe 2019). Additionally, the presence of tandem repeats within the control region was assessed using Tandem Repeats Finder (Benson, 1999). A consensus circular mitogenome figure was generated with Organellar Genome Draw “OGDRAW” version 1.3.1 (Greiner et al. 2019) and edited using Inkscape macOS version 1.4 (Inkscape Project, 2020).

### Bayesian phylogenetic analysis

A Bayesian phylogenetic analysis was conducted using a concatenated matrix of the 13 PCGs from 21 mitogenomes of closely related Myliobatiformes species across 8 families: Aetobatidae, Dasyatidae, Gymnuridae, Rhinopteridae, Mobulidae, Myliobatidae, Plesiobatidae, and Urolophidae. *Pristis pectinata* and *P. pristis* (Rhinopristiformes: Pristidae) were used as outgroups. Nucleotide sequences from each PCG were individually aligned using MEGA 7 (Kumar et al. 2016) and then concatenated using SeaView version 5.0.4 (Guoy et al. 2021). Substitution saturation in single codon positions from each PCG was analyzed using DAMBE 5 (Xia 2013). The best-fit model of evolution for each PCG was determined using jModelTest 2 (Darriba et al. 2012) under the corrected Bayesian Information Criterion (BICc). The third codon positions of three PCGs (ATP8, ND3, and ND6) exhibited clear signs of saturation; thus, the Bayesian analysis was performed with the data partitioned by gene and by codon position, excluding the third codon position of those genes. The phylogenetic tree was constructed using MrBayes 3.2.7 (Ronquist et al. 2012) as implemented on the CIPRES Science Gateway 3.3 server (Miller et al. 2010). This involved two independent runs of four Markov chains each, conducted for 20,000,000 generations, with a sampling frequency of 1,000. The first 25% of the sampled trees were discarded as burn-in. The final consensus phylogenetic tree was created using Figtree version 1.4.4 (http://tree.bio.ed.ac.uk/software/figtree/).

## Results and discussion

### Mitogenome assembly

The map to reference assembly analyses resulted in a total of 22,239 unmerged and 16,553 merged reads mapped to the reference sequence. These assembling processes produced two identical consensus sequences, each 20,217 bp in length. The novel mitogenome sequence of *A. ocellatus* was deposited in the GenBank database under accession number BK072016. It is important to note Third Party Annotations (TPA) submitted to GenBank must retain the same organism name as that in the primary sequence source. In this case, the primary sequence source referenced (BioSample accession SAMN31811701, SRA accession SRS15852991) belongs to a Biosample identified as *A. narinari* by the original submitters. Consequently, the newly assembled mitogenome sequence BK072016 retains the scientific name *A. narinari* in the GenBank nucleotide database, although the results from this study identified it as *A. ocellatus*.

### Mitogenome characterization

The mitogenome of *A. ocellatus* is 20,217 bp in length, making it larger than that of other related species within the Myliobatiformes, which typically range from 17,000 to nearly19,000 bp (Whitney et al. 2023). However, the mitogenome of *A. ocellatus* is only slightly larger than that of *A. flagellum*, which is 20,201 bp in length. There are also two additional mitogenome sequences for *Aetobatus* available in GenBank: *A. narinari*, (15,714 bp; GenBank accession KX151649) and *A. ocellatus* (11,901 bp; GenBank accession JN184054). However, both of these mitogenomes are incomplete, making it impossible to compare their sizes with the one reported here. Notably, relatively large mitogenomes have also been documented in other Chondrichthyan species, such as *Gymnura altavela* (19,472 bp; GenBank NC_059938) and *Rhinochimaera pacifica* (24,889 bp; HM147141). The different mitogenome sizes observed among several finfish and elasmobranch species are primarily attributed to a high content of tandem repeats within the control region (Kousteni et al. 2021).

The control region of the mitogenome in *A. ocellatus* measures 4,503 bp, accounting for 22.3% of the entire mitogenome. An analysis of the nucleotide sequence in this control region identified 8 distinct repeat motifs, varying in length from 24 to 75 bp, with copy numbers ranging from 2 to 9 repeats. The determination of additional mitogenome sequences from more individuals of *A. ocellatus* and other *Aetobatus* species will enhance our understanding of the nucleotide composition in the control region among members from this genus. This may reveal potential evolutionary signatures within *Aetobatus*.

Overall, the mitogenome of *A. ocellatus* exhibits a typical organization found in fish mitogenomes (Winn et al. 2024), as detailed in Table 1 and illustrated in Fig. 1. It comprises 13 PCGs, two rRNA genes, 22 tRNA genes, and one putative control region. The overall nucleotide composition of the mitogenome is as follows: A: 31.4%, C: 27.1%, G: 15.5%, and T: 26%. Of the total genes, 28 (75.7%) are located on the positive strand, while 9 (24.3%) are found on the negative strand (see Table 1 and Fig. 1). The total length of all PCGs in the novel mitogenome of *A. ocellatus* is 11,428 bp, which accounts for 56.5% of the entire mitogenome. The sizes of these genes vary, ranging from 168 bp for ATP8 to 1,833 bp for NAD5.

**Table 1.** Gene annotation of the mitochondrial genome of *A. ocellatus*.

| Gene | Strand | Location |  | Size (bp) | Anticodon | START codon | STOP codon | No. of amino acids |
| --- | --- | --- | --- | --- | --- | --- | --- | --- |
|  |  | Start | End |  |  |  |  |  |
| Phe (F) | + | 1 | 69 | 69 | GAA | — | — | — |
| 12S rRNA | + | 71 | 1031 | 961 | — | — | — | — |
| Val (V) | + | 1032 | 1103 | 72 | UAC | — | — | — |
| 16S rRNA | + | 1104 | 2807 | 1704 | — | — | — | — |
| Leu (L) | + | 2808 | 2882 | 75 | UAA | — | — | — |
| ND1 | + | 2883 | 3857 | 975 | — | ATG | TAA | 324 |
| Ile (I) | + | 3859 | 3926 | 68 | GAU | — | — | — |
| Gln (Q) | - | 3928 | 3999 | 72 | UUG | — | — | — |
| Met (M) | + | 4000 | 4069 | 70 | CAU | — | — | — |
| ND2 | + | 4070 | 5115 | 1046 | — | ATG | TA- | 348 |
| Trp (W) | + | 5116 | 5184 | 69 | UCA | — | — | — |
| Ala (A) | - | 5186 | 5254 | 69 | UGC | — | — | — |
| Asn (N) | - | 5256 | 5328 | 73 | GUU | — | — | — |
| Cys (C) | - | 5362 | 5430 | 69 | GCA | — | — | — |
| Tyr (Y) | - | 5436 | 5503 | 68 | GUA | — | — | — |
| CO1 | + | 5510 | 7063 | 1554 | — | GTG | TAA | 517 |
| Ser (S) | - | 7068 | 7138 | 71 | UGA | — | — | — |
| Asp (D) | + | 7139 | 7209 | 71 | GUC | — | — | — |
| CO2 | + | 7213 | 7903 | 691 | — | ATG | T-- | 230 |
| Lys (K) | + | 7904 | 7978 | 75 | UUU | — | — | — |
| ATP8 | + | 7980 | 8147 | 168 | — | ATG | TAA | 55 |
| ATP6 | + | 8138 | 8820 | 683 | — | ATG | TA- | 227 |
| CO3 | + | 8821 | 9606 | 786 | — | ATG | TAG | 261 |
| Gly (G) | + | 9615 | 9686 | 72 | UCC | — | — | — |
| ND3 | + | 9688 | 10036 | 349 | — | ATG | T-- | 116 |
| Arg (R) | + | 10037 | 10110 | 74 | UCG | — | — | — |
| ND4L | + | 10111 | 10407 | 297 | — | ATG | TAA | 98 |
| ND4 | + | 10401 | 11781 | 1381 | — | ATG | T-- | 460 |
| His (H) | + | 11782 | 11850 | 69 | GUG | — | — | — |
| Ser (S) | + | 11852 | 11920 | 69 | GCU | — | — | — |
| Leu (L) | + | 11923 | 11994 | 72 | UAG | — | — | — |
| ND5 | + | 11995 | 13827 | 1833 | — | ATG | TAA | 610 |
| ND6 | - | 13824 | 14345 | 522 | — | ATG | TAG | 173 |
| Glu (E) | - | 14347 | 14415 | 69 | UUC | — | — | — |
| CYTB | + | 14418 | 15560 | 1143 | — | ATG | TAA | 380 |
| Thr (T) | + | 15568 | 15641 | 74 | UGU | — | — | — |
| Pro (P) | - | 15646 | 15714 | 69 | UGG | — | — | — |
| Control region | + | 15715 | 20217 | 4503 | — | — | — | — |

**Fig. 1.**
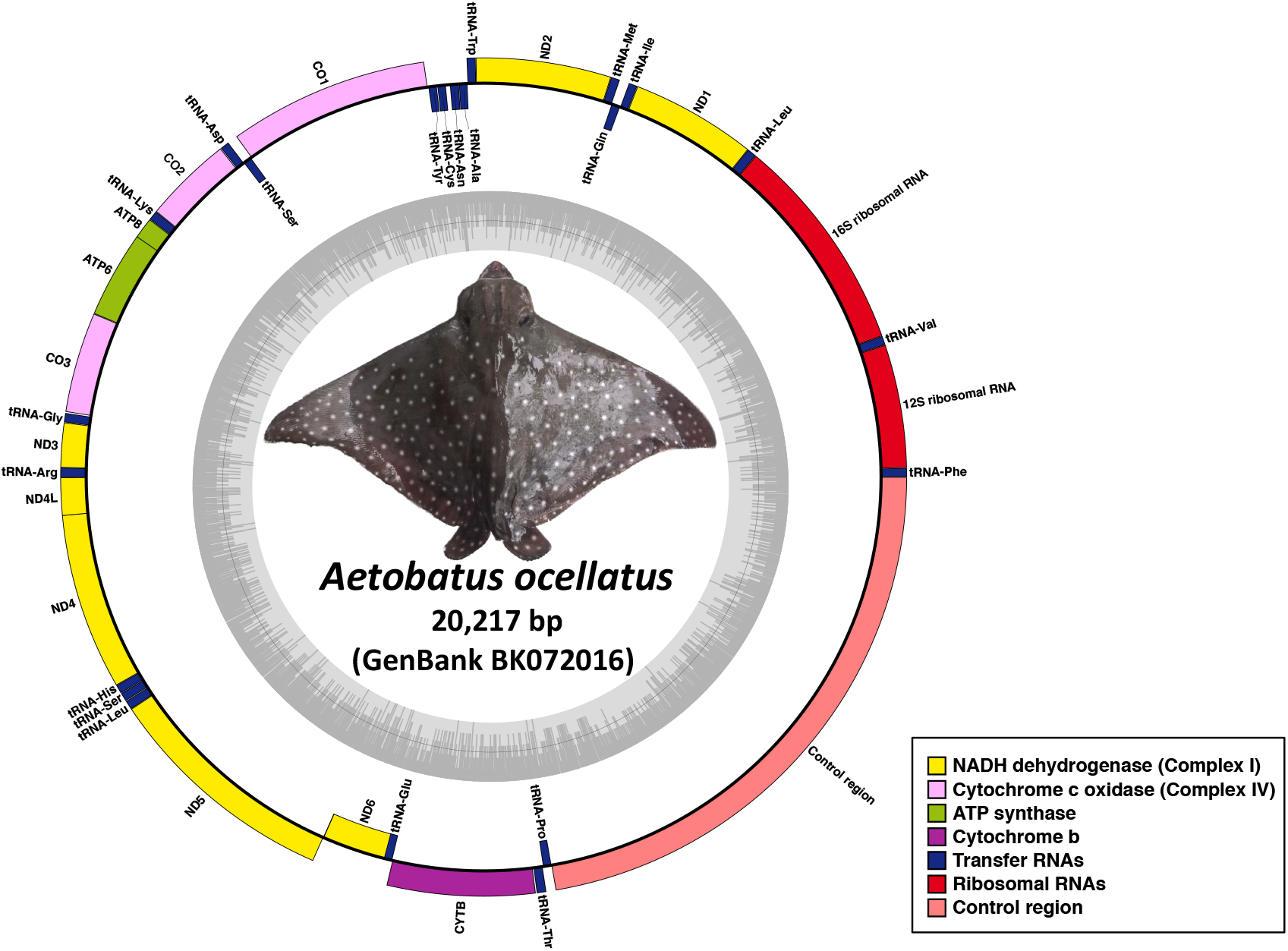
Circular representation of the mitochondrial genome of *Aetobatus ocellatus*. The specimen shown in the photograph (voucher USNM 435544, BioSample accession SAMN31811701) is the actual holotype from which the NGS genomic reads (SRA accession SRS15852991) were amplified by the original submitters. The photograph was taken by and reproduced with the permission from the Smithsonian Institution, NMNH, Division of Fishes, under the terms of a CC0 1.0 license (Orrell, 2025). In the diagram, genes located in inner circle (negative strand) are transcribed in a clockwise sense, while those on the outer circle (positive strand) are transcribed counterclockwise. The innermost circle of the GC content graph indicates a 50% threshold, with darker and lighter grey shades representing GC and AT content, respectively. Genes are represented by colored bars: ND dehydrogenase (yellow), cytochrome c oxidase (pink), ATP synthase (green), cytochrome b (purple), transfer RNA (blue), and ribosomal RNA (red). The control region is illustrated with a coral red bar.

All PCGs use the typical ATG start codon, except for the COI gene, which uses the GTG start codon. The standard stop codon TAA was detected in six PCGs: ATP8, CO1, CYTB, ND1, ND4L, and ND5. Five PCGs (ATP6, CO2, ND2, ND3, and ND4) use incomplete stop codons (TA- or T--), which are completed through a post-transcriptional polyadenylation process (Nagaike et al. 2005). Another standard stop codon, TAG, was observed in only two PCGs: CO3 and ND6.

### Phylogenetic analysis

The concatenated data from the 13 PCGs resulted in a final matrix comprising 11,468 nucleotides across 21 mitogenomes of Myliobatiformes, plus two additional mitogenomes of Rhinopristiformes that were used as outgroups. Fig. 2 presents a Bayesian phylogenetic tree where all samples are grouped into eight monophyletic family clades that exhibit high to strong nodal support. The overall structure of the phylogenetic tree revealed four distinct groups. Group I is situated at the top of the tree and includes the families Mobulidae and Rhinopteridae. Group II consists of the families Aetobatidae and Myliobatidae, while Group III is composed of three families: Gymnuridae, Urolophidae, and Plesiobatidae. At the base of the tree, Group IV contains the family Dasyatidae, which is sister to all other families within Myliobatiformes. These findings are consistent with those of previous related phylogenetic studies (Naylor et al. 2012; Lim et al. 2015).

**Fig. 2.**
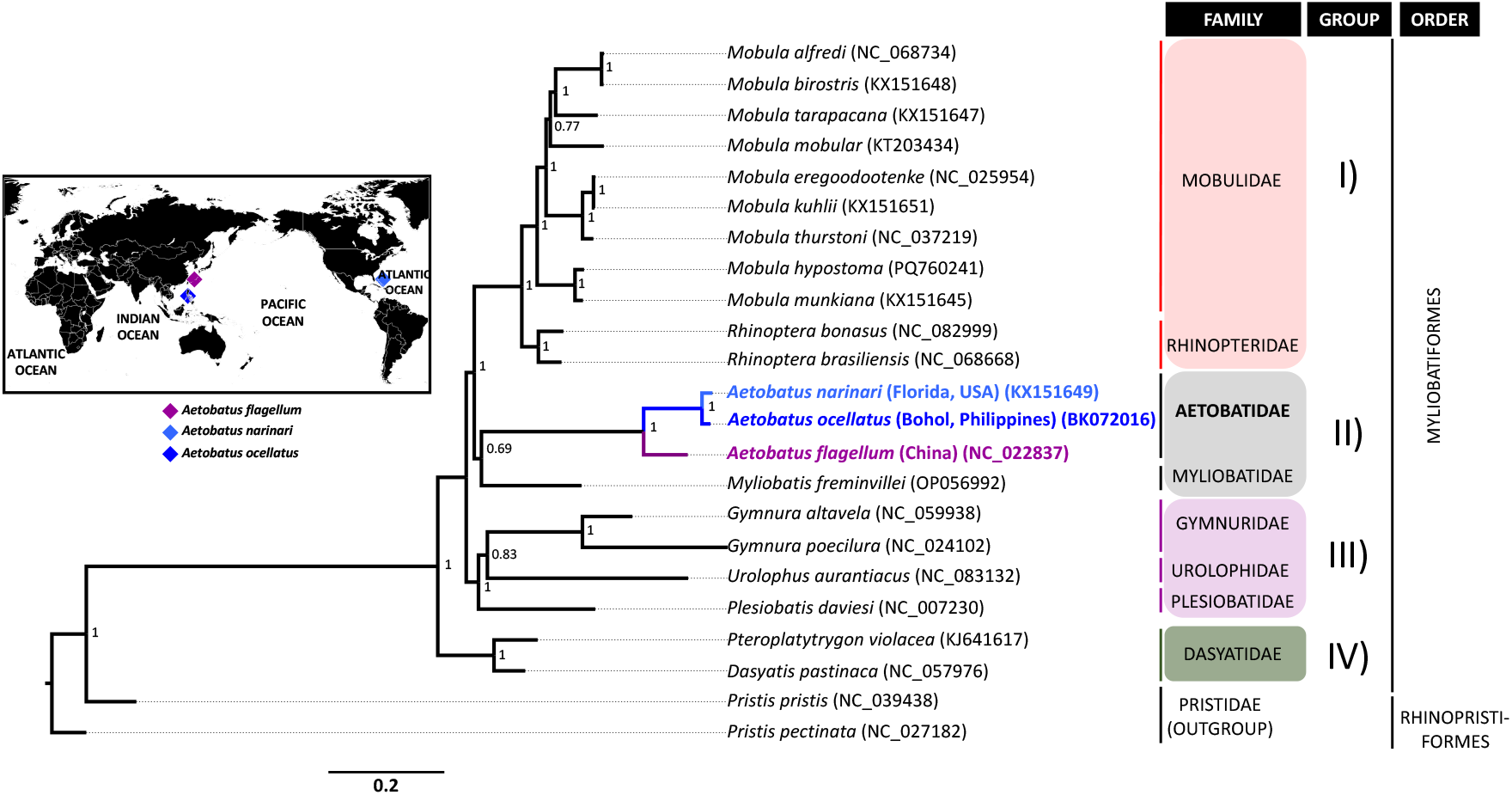
Bayesian phylogenetic tree inferred from 21 concatenated mitochondrial protein-coding genes from 21 Myliobatiformes species, representing 8 families: Aetobatidae, Dasyatidae, Gymnuridae, Mobulidae, Myliobatidae, Plesiobatidae, Rhinopteridae, and Urolophidae. The sequence matrix used for the phylogenetic analysis comprised 11,468 nt, focusing on unambiguously aligned regions from the first, second, and third codon positions. The position of *Aetobatus ocellatus* is highlighted in blue. The inset map illustrates the geographic origins of the three analyzed *Aetobatus* species. *Pristis pectinata* and *P. pristis* (Rhinopristiformes: Pristidae) were included as outgroups. Families that exhibited sister relationships are indicated in colored panels labeled with Roman numerals. Posterior probabilities are shown at each node, and GenBank accession numbers for each species are provided in parentheses.

The Aetobatidae clade was recovered as a well separated monophyletic group with robust statistical support. This finding reaffirms its resurrected family status, which was previously confused with Myliobatidae and Mobulidae (White and Naylor 2016).Within the Aetobatidae clade, *A. narinari* and *A. ocellatus* were clustered together in a discrete subclade, with their analyzed sequences diverging by 338 variable sites, reflecting a genetic divergence of 3.03%. These results suggest a relatively low divergence age, which has been previously estimated at 2 to 5 mya (Sales et al. 2019). The observed genetic distance between *A. ocellatus* and *A. narinari* falls within the range of interspecific genetic divergence rates that are suitable for species delineation found in related myliobatiform species (Bustamante et al. 2016; White et al. 2018).

Furthermore, the genetic evidence distinguishing *A. ocellatus* from *A. narinari* is also supported by the geographic origins of the analyzed samples. The genetic material used in this study (BioSample SAMN31811701) was collected from a specimen identified as *A. narinari* collected in the Indo-West Pacific region (Bohol Island, Philippines). In contrast, the mitogenome of *A. narinari* included in this analysis was obtained from the Atlantic Ocean (Florida, U.S.A.). The phylogenetic results obtained herein indicate that BioSample SAMN31811701 actually belongs to *A. ocellatus*. Overall, the findings of this research support the species status and classification of *A. ocellatus* as an Indo-West Pacific species (White et al. 2010), while *A. narinari* is restricted to the Atlantic region (Sales et al. 2019).

## Conclusion

This study used NGS data from GenBank to assemble the first mitochondrial genome of the endangered spotted eagle ray, *Aetobatus ocellatus*. The mitogenome was characterized and subjected to a comprehensive phylogenetic analysis alongside related species within the Myliobatiformes order. The phylogenetic outcome reinforced the classification of Aetobatidae as a valid family and confirmed *A. ocellatus* as a recognized species. Before this study, the validation of *A. ocellatus* was supported by taxonomic and phylogenetic analyses based on a single mitochondrial gene (White et al. 2010). The results of this study significantly contribute to a deeper understanding of the complex taxonomic and phylogenetic relationships within Aetobatidae. These findings could aid conservation efforts and taxonomic clarity for the vulnerable species from this genus, whose taxonomic classification is still under debate.

## Data availability

The mitogenome nucleotide sequence data reported is available in the Third Party Annotation Section of the DDBJ/ENA/GenBank databases under the accession number TPA: BK072016.

